# Extending the proteomic characterization of *Candida albicans* exposed to stress and apoptotic inducers through data-independent acquisition mass spectrometry

**DOI:** 10.1101/2020.12.20.423720

**Authors:** Ahinara Amador-García, Inés Zapico, Johan Malmström, Lucía Monteoliva, Concha Gil

**Affiliations:** Department of Microbiology and Parasitology, Faculty of Pharmacy, Ramon y Cajal Health Research Institute (IRYCIS), Complutense University of Madrid (UCM), Madrid, Spain; Proteomics Unit, Complutense University of Madrid, Spain; Division of Infection Medicine, Department of clinical Sciences, Lund University, Lund, Sweden

**Keywords:** *C. albicans*, oxidative stress, acetic acid, proteomics, data-independent acquisition, selected reaction monitoring, apoptosis, proteasome

## Abstract

*Candida albicans* is a commensal fungus that causes systemic infections in immunosuppressed patients. In order to deal with the changing environment during commensalism or infection, *C. albicans* must reprogram its proteome. Characterizing the stress-induced changes in the proteome that *C. albicans* uses to survive should be very useful in the development of new antifungal drugs. We studied the *C. albicans* global proteome after exposure to hydrogen peroxide (H_2_O_2_) and acetic acid (AA), using a DIA-MS strategy. More than 2000 *C. albicans* proteins were quantified using an ion library previously constructed using DDA-MS. *C. albicans* responded to treatment with H_2_O_2_ with an increase in the abundance of many proteins involved in the oxidative stress response, protein folding and proteasome-dependent catabolism, which led to an increased proteasome activity. The data revealed a previously unknown key role for Prn1, a protein similar to pirins, in the oxidative stress response. Treatment with AA resulted in a general decrease in the abundance of proteins involved in amino acid biosynthesis, protein folding, and rRNA processing. Almost all proteasome proteins declined, as did proteasome activity. Apoptosis was observed after treatment with H_2_O_2_, but not AA. A targeted proteomic study of 32 proteins related to apoptosis in yeast supported the results found by DIA-MS and allowed the creation of an efficient method to quantify relevant proteins after treatment with stressors (H_2_O_2_, AA, and amphotericin B). This approach also uncovered a main role for Oye32, an oxidoreductase, suggesting this protein as a possible apoptotic marker common to many stressors.

**IMPORTANCE:** Fungal infections are a worldwide health problem especially in immunocompromised patients and patients with chronic disorders. Invasive candidiasis, mainly caused by *C. albicans*, are among the most common fungal diseases. Despite the existence of treatments to combat candidiasis the spectra of drugs available are limited. For the discovery of new drug targets is essential to know the pathogen response to different stress conditions. Our study provides a global vision of proteomic remodeling in *C. albicans* after exposure to different agents such as hydrogen peroxide, acetic acid and amphotericin B that can cause apoptotic cell death. This results revealed the significance of many proteins related to oxidative stress response and proteasome activity among others. Of note, the discovery of Prn1 as a key protein in the defence against oxidative stress as well the increase in the abundance of Oye32 protein when apoptotic process occurred point out them as possible drug targets.

## INTRODUCTION

*C. albicans* is a common opportunistic fungus in the human microbiota that can cause severe infections in immunocompromised hosts. Candidiasis, caused mainly by *C. albicans*, ranges from local mucosal to systemic infections and has a noteworthy clinical impact on morbidity and mortality in intensive-care-unit patients (1). In spite of current antifungal therapies, it is estimated that invasive candidiasis causes 50,000 deaths worldwide every year (2). This is partially explained by the increase in the number of high-risk hosts or a late/deficient diagnosis, but is also due to emerging resistance to antifungal drugs.

Among other pathogenicity mechanisms, *C. albicans* has the ability to respond and adapt to different host microenvironments (3), including a wide pH range and the anti-microbial oxidative burst originated by innate immune cells. *C. albicans* has developed several antioxidant mechanism including catalase, superoxide dismutases (SODs), and gluthatione and thioredoxin systems that enzymatically detoxify O_2_ radicals such as hydrogen peroxide (4). As a member of the gut microbiota, *C. albicans* also has to cope with metabolites, including weak organic acids such as acetic acid, produced by other microorganisms (5). Both hydrogen peroxide and acetic acid have been described as inducers of regulated cell death in *C. albicans* (6-8). For this reason, the study of the response of *C. albicans* to these agents is a promising alternative strategy against this pathogen. Regulated cell death in *C. albicans* upon exposure to many agents, including antifungals and plant extracts, has been widely demonstrated (9-13). Many studies in *C. albicans* and *Saccharomyces cerevisiae* have revealed the participation of several proteins in this process, mainly from mitochondria and the Ras pathway as well as metacaspase I and its substrates (14-16). Although the terms *programmed cell death* and *apoptosis* are sometimes used interchangeably to describe regulated cell-death processes, their precise meanings can be distinguished using the recommended guidelines for yeast cell-death nomenclature (17). DNA fragmentation, an increase in caspase-like enzymatic activity, and reactive oxygen species production or phosphatidylserine (PS) exposure have been widely considered as apoptotic markers (18, 19). However, only PS exposure is currently accepted as an apoptotic marker in yeast, and *programmed cell death* applies to cell death triggered under physiological scenarios such as aging. The term *regulated cell death* includes both apoptosis and programmed cell death as organized cell death processes promoted by external or internal stresses.

In order to deal with the changing environment during commensalism or infection, *C. albicans* must reprogram its proteome by expressing or repressing certain proteins. A better characterization of these changes in response to stress inducers is important for a deep understanding of *C. albicans* survival strategies, which would be very useful in the development of new antifungal therapies to combat the infection.

With this purpose, we used a data-independent acquisition (DIA) proteomic approach to identify global changes in the abundance of *C. albicans* proteins in response to oxidative and acetic acid stresses. This strategy for global proteomic studies is an improvement compared with the traditional data-dependent (DDA) approach, conferring better quantitative accuracy and reproducibility and enlarging the number of quantifiable peptides (20). Taking advantage of targeted proteomics as well, we used a selected reaction monitoring (SRM) method to monitor the abundance of proteins related to regulated cell death (21). This SRM method allowed the straightforward quantitation of key proteins involved in regulated cell death after exposure to the stressors hydrogen peroxide, acetic acid, and amphotericin B, an antifungal previously described as an apoptotic inducer (16, 22).

## RESULTS

### Effects of hydrogen peroxide and acetic acid on viability, physiological response, and cell death in *C. albicans*

The main goal of this study was to evaluate proteomic changes in cells exposed to different concentrations of hydrogen peroxide and acetic acid in order to reveal the main processes involved in stress responses. With the aim of calculating the damage produced by these agents on growth and cellular viability, optical density and colony formation were measured after treatment. The results show that the growth and viability of cells exposed to hydrogen peroxide were compromised in a dose-dependent manner. The final optical densities of the cell cultures after treatment with 5 mM and 10 mM hydrogen peroxide were, respectively, 2.5 and 3.4 times lower than those of the control cells, and only 35% and 13%, respectively, of the cells were viable. In contrast, acetic acid at 40 mM and 60 mM induced cell growth arrest but did not lower cell viability, as indicated by the recovery of cell growth in colonies after the treatment was stopped (Fig. 1). Despite the effects observed, the loss of cell membrane integrity as measured by propidium iodide (PI) staining was in all cases below 1%.

**Figure 1.**
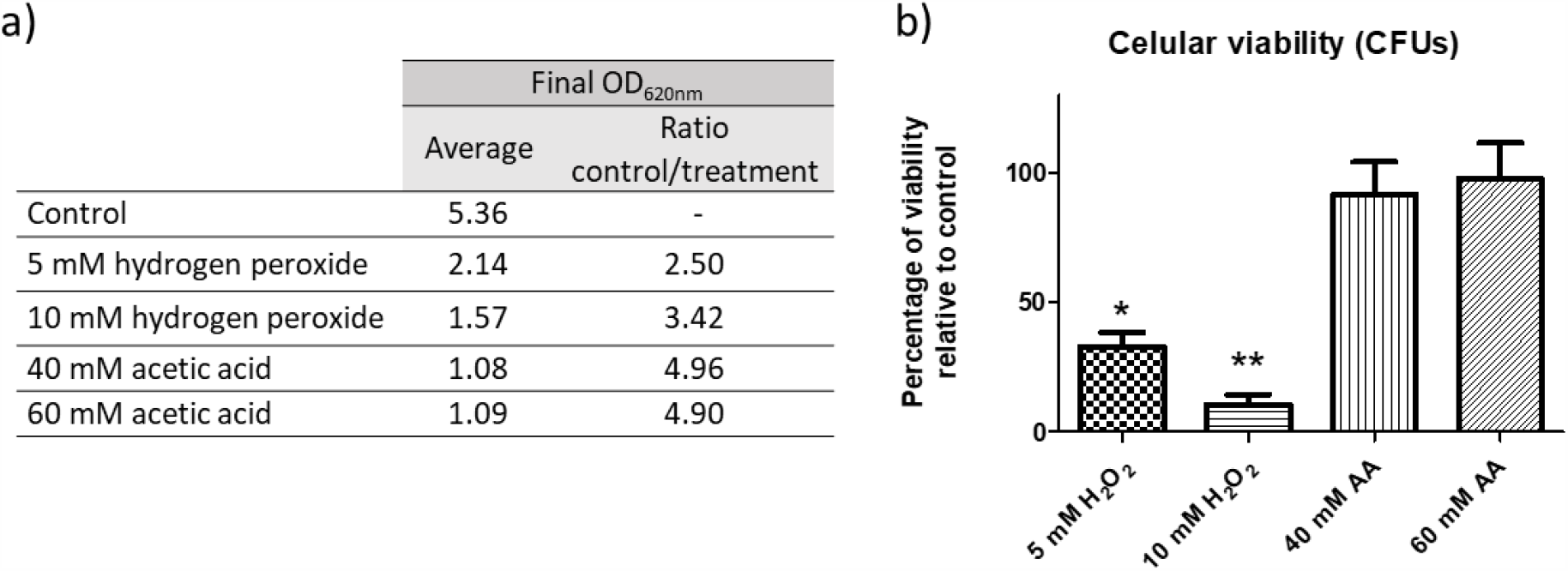
Effects on *C. albicans* growth after exposure to hydrogen peroxide or acetic acid. a) Optical density reached after treatment for 200 min with the stress inducers cited. b) Percentage of viability in *C. albicans* cells treated with the agents compared to control samples. A significant change is indicated as *pvalue<0.05 or **pvalue<0.01* (paired t-test). All results represent the average of at least three biological replicates.

In order to evaluate the physiological response of *C. albicans* to the agents, we measured ROS production and caspase-like enzymatic activity (17). Both increased after all the treatments studied, most remarkably at the highest concentrations used (Fig. 2A). The ROS production in 20% and 34% of cells exposed to 10 mM hydrogen peroxide and 60 mM acetic acid, respectively, confirmed the oxidative stress promoted by the agents. Moreover, a moderate, non-significant increase in caspase-like enzymatic activity demonstrated an active response of the cells to the compounds (Fig. 2B). Apoptosis evaluated by PS externalization was promoted by both concentrations of hydrogen peroxide, reaching significance after exposure to 10 mM H_2_O_2_, which resulted in up to 40% of cells becoming apoptotic (Fig. 2C). Nevertheless, the absence of a significant increase in this marker in cells exposed to acetic acid ruled out an apoptotic response under the conditions used.

**Figure 2.**
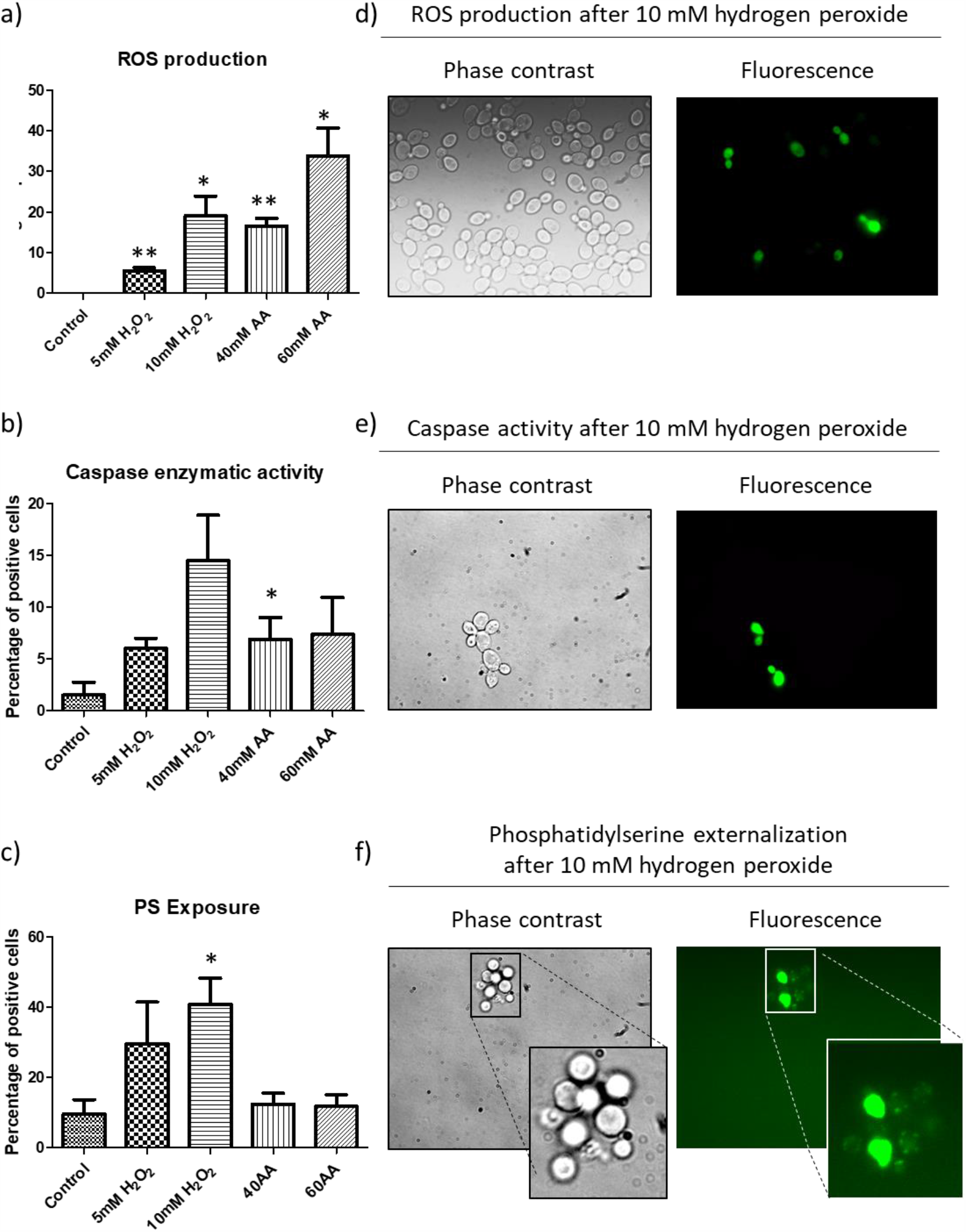
Apoptotic markers analyzed in *C. albicans*. Graphics show the percentage of positive cells for ROS (a), caspase like enzymatic activity (b), and PS exposure (c). Results represent the mean of at least three biological replicates. Cells were counted on a fluorescence microscope and significant changes are indicated (paired t-test). Representative fluorescence microscopy images from cells treated with hydrogen peroxide showing ROS (d), caspase activity (e) and PS exposure (f).

### Quantitative profiling of the *C. albicans* proteome after exposure to stressors

To acquire a representative picture of changes in the *C. albicans* global proteome after exposure to hydrogen peroxide and acetic acid, we used DIA-MS. This approach allowed us to quantify more than 2000 *C. albicans* proteins in the four conditions tested by using the ion library previously constructed using DDA-MS. It comprises information on almost 46.5% of the *C. albicans* proteome (unpublished data). Statistical analysis revealed a remarkable remodeling of the proteome, involving changes in the abundance of hundreds of proteins under each condition (Fig. 3). The proteomic responses observed after hydrogen peroxide or acetic acid treatment differed greatly. While the treatment with hydrogen peroxide predominantly promoted an increase in the abundance of a large number of proteins, the exposure to acetic acid was characterized by a profound decrease in the abundance of many proteins (Fig. 3). These dissimilar patterns, which are clearly seen in the volcano plots, are evidence of specific responses of this fungus to the two stress inducers studied.

**Figure 3.**
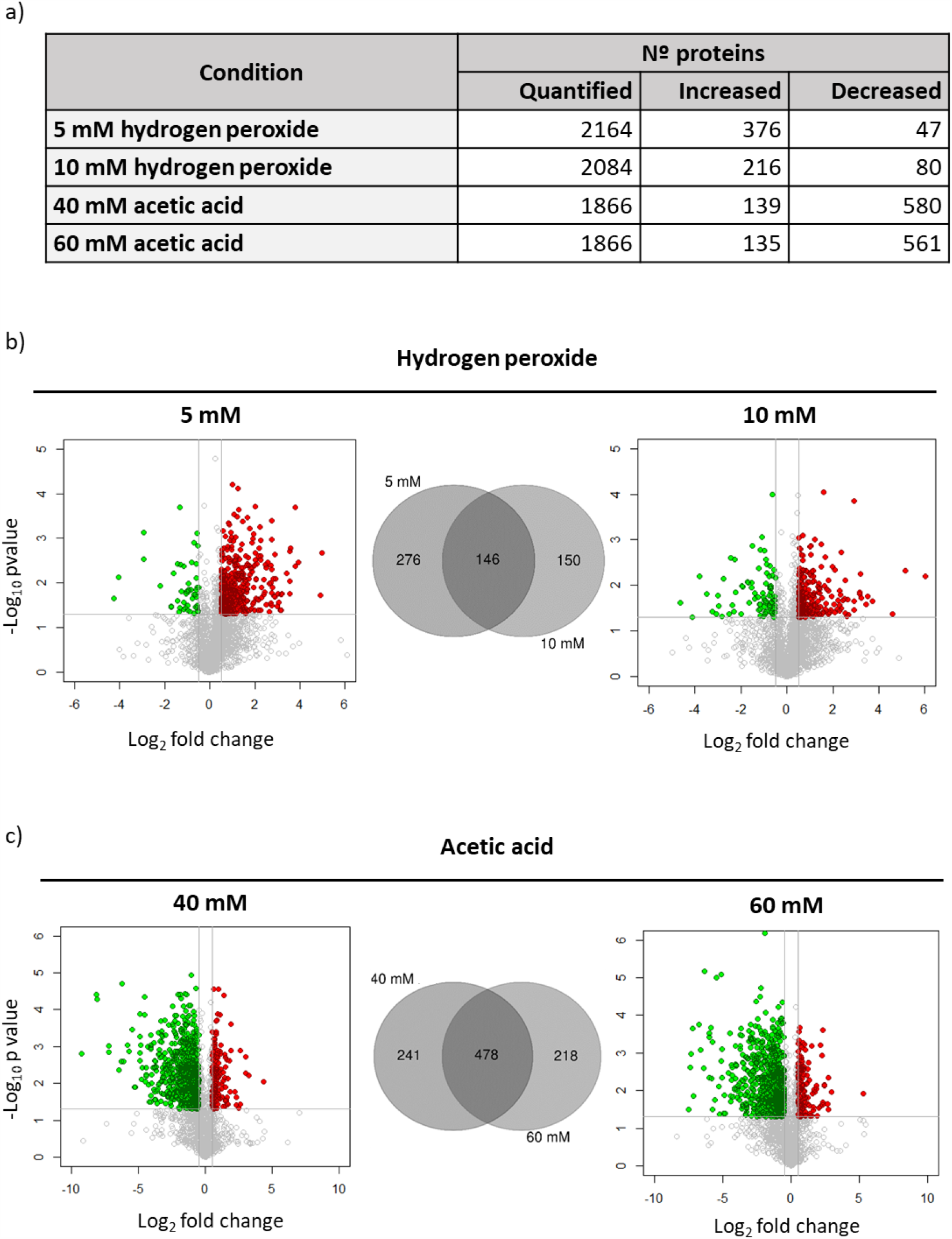
Results from DIA quantitation after exposure to hydrogen peroxide and acetic acid. a) Number of quantified proteins and proteins with changes in their abundance in each condition. Volcano plots representing results from DIA analysis comparing cells with or without treatment with hydrogen peroxide (b) or acetic acid (c). Changes in the abundance of proteins are represented in red or green according to their significant increase (-log_10_pval>1.3; log_2_ fold change>0.5) or decrease (-log_10_pva>1.3; log_2_ fold change<-0.5). Venn diagram showing the number of proteins with common or specific changes at the two concentrations of hydrogen peroxide (b) or acetic acid (c) tested.

To further understand the effects of the agents on biological processes, we use GO enrichment analysis to characterize proteins whose abundance changed significantly. Significant processes (p<0.05) containing the highest number of proteins from GO enrichment analysis were represented in heatmaps jointly for all treatments. This revealed an increase in key proteins related to the oxidative stress response, proteasome-dependent catabolism, and protein folding after treatment with hydrogen peroxide (Fig. 4 and Table S1). A total of 38 and 25 proteins involved in the oxidative stress response were more abundant after treatment with 5 mM and 10 mM hydrogen peroxide, respectively. Among them were essential members of the main detoxification system in *C. albicans* such as Cat1, the superoxide dismutases Sod1 and Sod2, glutaredoxin Ttr1, and the thioredoxins Tsa1, Trx1, and Trr1 (4, 23). Another relevant process unmasked by the GO analysis was cellular catabolism, with notable increases in the levels of 101 and 34 proteins after treatments with 5 mM and 10 mM hydrogen peroxide, respectively. A total of 40 and 20 proteins involved in catabolism mediated by the proteasome were increased in abundance after treatment at the concentrations mentioned, with more proteins increasing in abundance in response to 5 mM than to 10 mM hydrogen peroxide.

**Figure 4.**
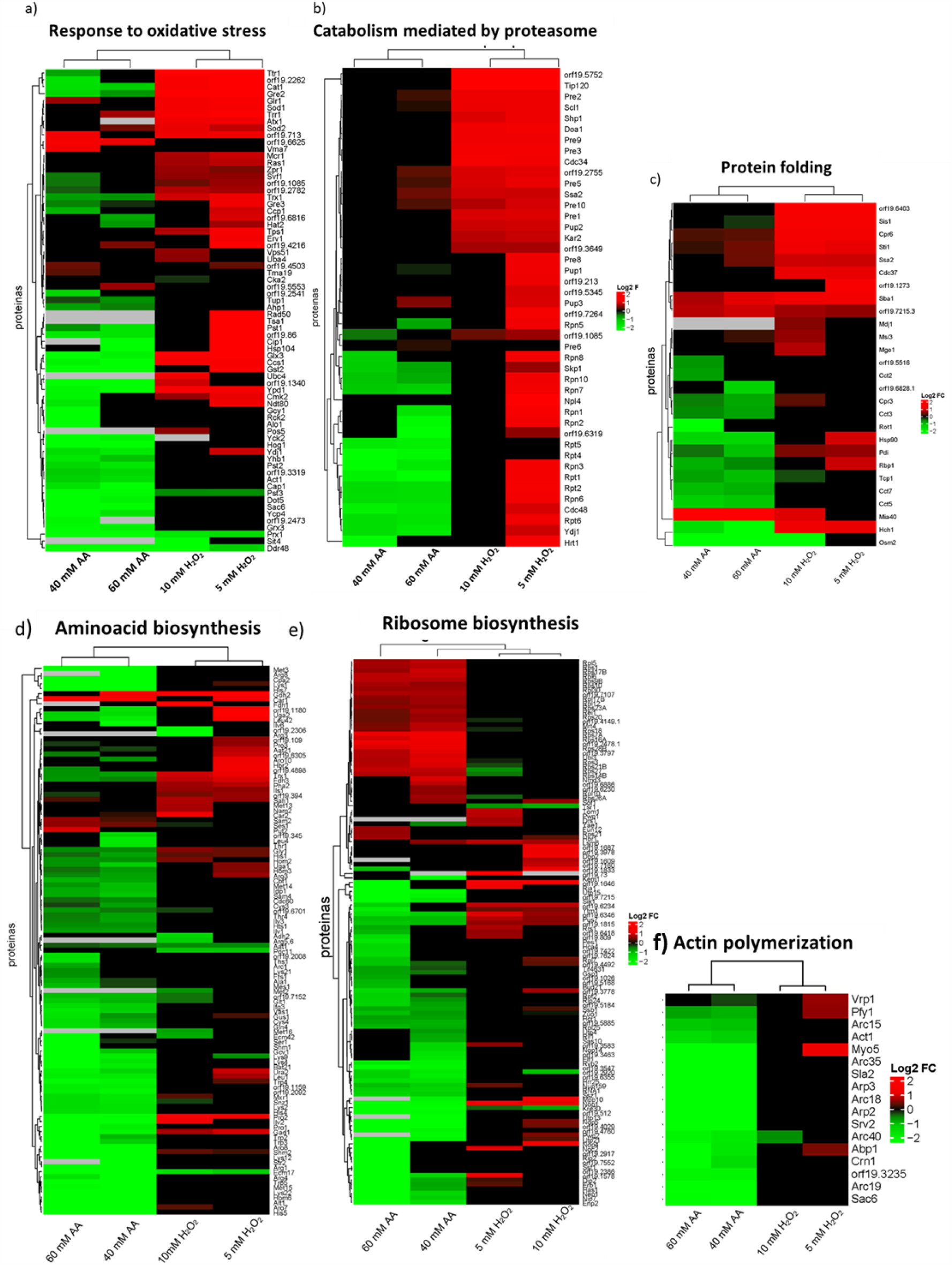
Heatmaps containing proteins that changed their abundance after treatments belonging to relevant biological processes. (a) Response to oxidative stress, (b) catabolism mediated by proteasome, (c) protein folding, (d) aminoacid biosynthesis, (e) ribosome biosynthesis or (f) actin polymeration. In red or green are represented proteins that increased or decreased their abundance after treatment with hydrogen peroxide (right columns) or acetic acid (left columns).

Although the heatmaps in Fig. 4 show mostly increases in most significant processes after treatments with hydrogen peroxide, other interesting groups of proteins from the respiratory chain (Cox4, Cox5, Cox6, Cox8, Cox9, and Qcr7) and cell wall (Ecm33, Pga4, Pga52, Sun41, and Tos1) had decreased levels in these conditions (Table S2).

The three biological processes with increased protein levels after treatment with hydrogen peroxide showed a completely opposite response when cells were treated with acetic acid, where a general decrease was observed (Fig. 4). These included proteins involved in oxidative response to stress, but many heat shock proteins were also less abundant after acetic acid treatment, so that the total number of proteins included in the stress response was very large. In addition, other biological processes were also characterized by a general decrease in protein levels. This up to 100 proteins participating in the biosynthesis of most amino acids were diminished in abundance after acetic acid exposure, indicating almost complete repression of this process (Fig. 4 and Table S3). Also, the actin cytoskeleton organization GO process was enriched, including proteins from the Arp2/3 family and proteins involved in actin folding (the CCT chaperone complex), that decreased their abundance after acetic acid exposure (Fig. 4). The levels of more than 30 proteins from the small and large subunits of the ribosome (in the Rps and Rpl families) increased. Conversely, the levels of 48 proteins involved in rRNA processing (e.g., Nop5, Rrs1, Utp13, Utp15, Utp21) declined.

### Opposite impacts on proteasome activity after hydrogen peroxide and acetic acid treatment in *C. albicans*

Our proteomic approach demonstrated that hydrogen peroxide and acetic acid treatments had opposite effects on the abundance of proteasome proteins. The proteasome is responsible for the specific degradation of abnormal, short-lived, and regulatory proteins, and comprises a central catalytic component (20S) with major three major proteolytic activities: chymotrypsin-like, trypsin-like, and peptidyl-glutamyl peptide hydrolyzing activities, and a regulatory particle (19S) conferring ATP and ubiquitin dependence on protein degradation (24).

Our data showed an increase in the expression levels of proteasome subunits after hydrogen peroxide treatment. This was more noticeable after 5 mM hydrogen peroxide treatment, with increased levels of 21 out of 25 quantified proteasome proteins representing both the regulatory and the central core particle (Fig. 5A). After 10 mM hydrogen peroxide treatment, only seven proteins from the catalytic particle were more abundant (Pre1, Pre2, Pre3, Pre5, Pre9, Pre10, and Pup2). In contrast, a dramatic decrease in proteasome protein levels occurred after 40 mM and 60 mM acetic acid treatment. Surprisingly, this affected only proteins from the regulatory particle (Rpn 1, Rpn2, Rpn3, Rpn5, Rpn6, Rpn7, Rpn8, Rpt1, Rpt2, Rpt4, Rpt5 and Rpt6) (Fig. 5A).

**Figure 5.**
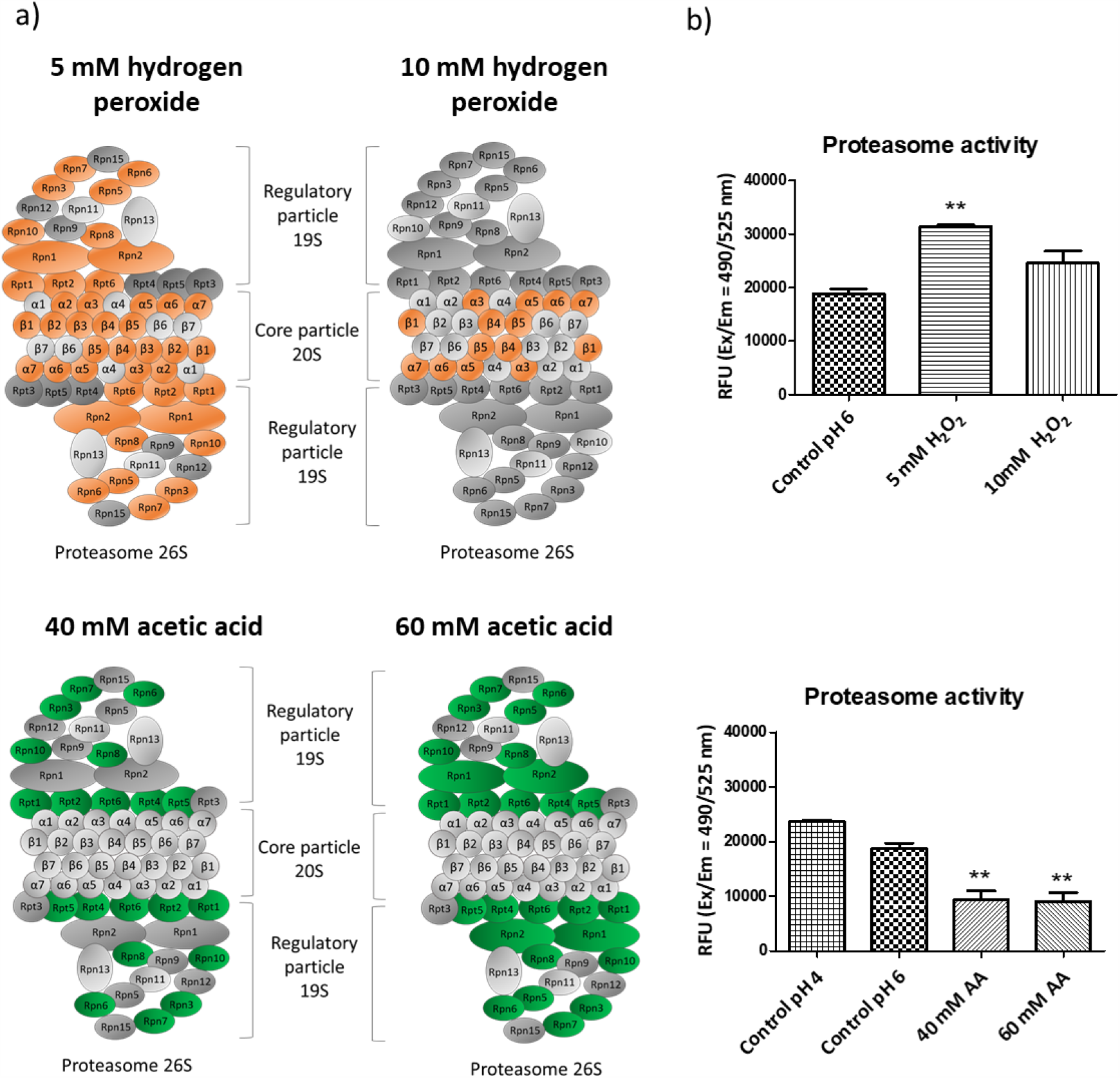
Changes in the abundance of proteins from the proteasome after the exposure to acetic acid or hydrogen peroxide. (a) In orange or green are highlighted proteins that increased or decreased significantly. (b) Results from the measurement of chymotrypsin-protease activity of the proteasome in RFU (relative fluorescence units).

In order to correlate these changes in the abundance of proteasome proteins to differences in proteasome activity, the chymotrypsin-like protease activity associated with the proteasome complex was measured. The results entirely corroborate our proteomic findings. The exposure to hydrogen peroxide increased proteasome activity while the acetic acid treatment caused a 50% loss of activity. To ensure that this effect was not related to the drop in pH caused by acetic acid treatment, a control sample acidified by HCl was also tested and the decrease in proteasome activity was not observed. In addition, the dose-response effect observed by proteomics in relation to the two concentrations of hydrogen peroxide was also correlated with proteasome activity, which increased by 67% after 5 mM hydrogen peroxide treatment but by only 31% after 10 mM hydrogen peroxide treatment, with respect to the control (Fig. 5B).

### Proteomic analysis unmasks proteins highly relevant to oxidative stress

Focusing on the top ten proteins with the greatest increase in abundance after treatment with both concentrations of hydrogen peroxide (Table 1) we found proteins associated with the proteasome (Hsm3, orf19.5752, and Tip120) and response to oxidative stress (Gre2, Oye2, and Oye23). Also among those with the most prominent increases in expression were proteins involved in DNA repair (Hsm3), DNA replication (Mcm2), and iron and copper metabolism (orf19.2067 and Cup1).

**Table 1.** List with the top 10 proteins showing the highest increase in relative abundance after treatment with hydrogen peroxide.

| Identifier | Protein | Description | Biological process | Hydrogen peroxide |  |  |  |
| --- | --- | --- | --- | --- | --- | --- | --- |
|  |  |  |  | 5 mM |  | 10 mM |  |
|  |  |  |  | Ratio log<sub>2</sub> | p value | Ratio log<sub>2</sub> | p value |
| orf19.3150 | Gre2 | Putative reductase | Response to oxidative and osmotic stress | 4.97 | 0.00 | 4.60 | 0.04 |
| orf19.2467 | Prn1 | Protein with similarity to pirins | Unknown | 4.90 | 0.02 | 5.15 | 0.00 |
| orf19.6729 | Tip120 | Protein involved in regulation of SCF* | Proteasomal catabolic process | 3.58 | 0.00 | 3.16 | 0.02 |
| orf19.5752 | - | Protein involved in regulation of SCF* | Proteasomal catabolic process | 3.56 | 0.02 | 3.45 | 0.03 |
| orf19.3443 | Oye2 | NADPH dehydrogenase | Oxidation-reduction process. Apoptosis | 3.55 | 0.00 | 3.15 | 0.01 |
| orf19.3433 | Oye23 | NADPH dehydrogenase | Oxidation-reduction process | 3.54 | 0.01 | 3.23 | 0.02 |
| orf19.2067 | - | Mitochondrial iron metabolism | Iron ion binding | 3.40 | 0.01 | 3.19 | 0.01 |
| orf19.3940.1 | Cup1 | Metallothionein. Copper resistance | Copper ion binding | 3.20 | 0.02 | 2.93 | 0.03 |
| orf19.1331 | Hsm3 | Proteasome regulatory particle | Proteasome regulation and mismatch repair | 2.92 | 0.01 | 2.59 | 0.05 |
| orf19.4354 | Mcm2 | Phosphorylated protein of unknown function | DNA replication | 2.87 | 0.02 | 2.93 | 0.02 |

\*SCF: Skp, Cullin, F-box containing complex

The increase in the level of Prn1, a protein similar to pirins, whose function, biological process, and cellular component remain unknown in *C. albicans*, was particularly interesting. This protein exhibited the highest increase after treatment with 10 mM hydrogen peroxide and the second highest after treatment with 5 mM hydrogen peroxide. Furthermore, Prn1 was the only protein with a greater increase after treatment with 10 mM than with 5 mM hydrogen peroxide. This result suggests a main role for Prn1 in the response to oxidative stress. Analysis of predicted functional partners of this protein by String software showed networks with other key proteins in oxidative stress. In addition, most of them also increased their abundance in our experiments (Fig. 6A). To phenotypically validate this we analyzed the susceptibility of the mutant *prn1*Δ to hydrogen peroxide in comparison with wild-type strains. As shown in Fig. 6, the susceptibility of the *prn1*Δ mutant to 80 mM and 100 mM hydrogen peroxide was notably higher compared with the strain used in this work (SC5314) and the control strain from the Noble collection (SN250)(Fig. 6B).

**Figure 6.**
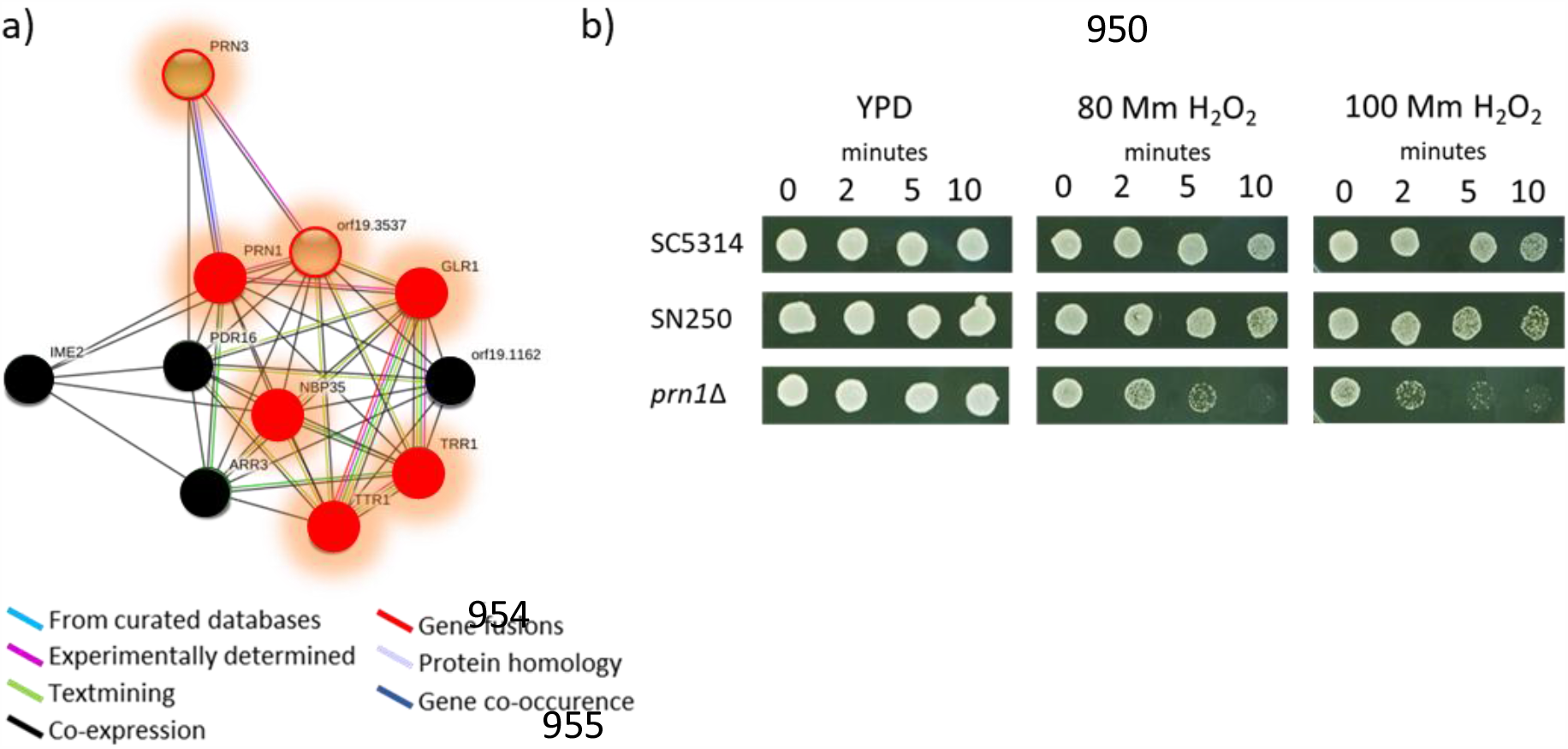
Prn1 protein in oxidative stress. A) Network showing predicted functional partners of Prn1 according to String software. In red, proteins that increased their abundance after exposure to hydrogen peroxide in this work. Proteins circled in red were only quantified after hydrogen peroxide exposure. In black, proteins that were not detected. B) Sensitivity to 80 and 100 mM hydrogen peroxide of the wild-type strains SC5314 and SN250 compared to *prn1Δ* mutant.

### The SRM method for quantitation of apoptotic proteins after exposure to stress inducers

Global proteomic results provide a panoramic picture of protein changes, but we were also very interested in specific proteins involved in regulated cell death. We chose targeted proteomics as the most useful and efficient means of analyzing them. We developed an SRM method to study these proteins. First, we selected 32 *C. albicans* proteins previously described (in *C. albicans* or *S. cerevisiae*) as relevant to yeast apoptosis involving mitochondria, the Ras pathway, or Mca1 (Table 2)(8, 14-16, 18, 25-54). SRM quantitation of 63 peptides and 462 transitions was performed on samples from *C. albicans* exposed to 5 mM or 10 mM hydrogen peroxide and 40 mM or 60 mM acetic acid (Table S4). From the 32 proteins included in the SRM method, 22 were quantified in at least one condition. Comparison of DIA and SRM quantitation showed that some proteins were only quantified by one proteomic approach.. The levels of proteins that were quantified in both the global DIA and the targeted SRM analysis were compared. In most cases, the two methods showed similar changes in protein levels (Table 3). The targeted approach revealed changes in the abundance of dozens of the selected proteins in most of the conditions tested, confirming the importance of these proteins upon exposure to these agents (Table 4). Results from the SRM quantitation showed a global pattern similar to that observed in the DIA analysis, with increased and decreased levels of many proteins selected after hydrogen peroxide and acetic acid exposure, respectively.

**Table 2.** Proteins involved in *C. albicans* or *S. cerevisiae* (*S*.*c*) apoptosis selected for SRM analysis. Proteins are related to regulated cell death by mitochondria, Ras/PKA/AMPc pathway, metacaspase 1 (Mca1) or other pathways.

| *C. albicans* | *S.c* | Description | Apoptosis<sup>a</sup> | References |  |
| --- | --- | --- | --- | --- | --- |
| Cpd1 | Aif1 | Mitochondrial apoptosis-inducing factor | PRO | (14, 15, 35, 38, 53) | Mitochondria |
| Cpr3 | Cpr3 | Putative peptidil-propyl cis-trans isomerase | PRO | (33, 35) |  |
| Cyc1 | Cyc1 | Cytochrome c | PRO | (15, 37, 40) |  |
| Dnm1 | Dnm1 | Putative dynamin-related GTPase | PRO | (15, 28) |  |
| Nde1 | Nde1 | Putative NADH dehydrogenase | PRO | (15, 35) |  |
| Nuc1 | Nuc1 | Major mitochondrial nuclease | PRO | (15, 38) |  |
| orf19.4423 | Ysp2 | Mitochondrial glucosyltransferase | PRO | (15, 43) |  |
| orf19.916 | Bxi1 | Protein involved in apoptosis | ANTI | (26, 27) |  |
| Oye2 | Oye2 | NAPDH dehydrogenase | ANTI | (29, 41) |  |
| Oye32 | Oye3 | NAD(P)H oxidoreductase | PRO | (41, 52) |  |
| Pet9 | Pet9 | Mitochondrial ADP/ATP carrier protein involved in ATP biosynthesis | PRO | (42) |  |
| Por1 | Por1 | Mitochondrial outer membrane porin | PRO/ANTI | (29, 34, 42, 54) |  |
| Rsm23 | Rsm23 | Mitochondrial ribosomal subunit | PRO | (25) |  |
| Sod2 | Sod2 | Mitochondrial superoxide dismutase | ANTI | (15) |  |
| Tma19 | Tma19 | Translation Machinery Associated | PRO | (15, 45) |  |
| Ymx6 | Nde1 | NADH dehydrogenase | PRO | (15, 32, 35) |  |
| Bcy1 | Bcy1 | Protein kinase A regulatory subunit | ANTI | (8) | Ras/PKA/AMPC |
| Cyr1 | Cyr1 | Class III adenylyl cyclase |  | (16) |  |
| Efg1 | Sok2 | bHLH transcription factor | ANTI | (49) |  |
| Ras1 | Ras2 | RAS signal transduction GTPase | PRO | (8, 14) |  |
| Tpk1 | Tpk1 | cAMP-dependent protein kinase catalytic subunit | PRO | (8) |  |
| Cdc48 | Cdc48 | Putative microsomal ATPase | ANTI | (14, 18, 31) | Mca1 and substrates |
| Mca1 | Mca1 | Putative metacaspase involved in apoptosis | PRO | (14, 31, 39) |  |
| Nma111 | Nma111 | Putative serine protease | PRO | (15, 29, 38, 50, 51) |  |
| orf19.643 | Bir1 | Ortholog have role in apoptosis | ANTI | (15, 29, 38, 51) |  |
| Tdh3 | Tdh3 | Glyceraldehyde-3-phosphate dehydrogenase | PRO | (46) |  |
| Wwm1 | Wwm1 | Protein of unknown function | PRO | (31, 47) |  |
| Cst20 | Ste20 | Protein kinase | PRO | (15) | Others |
| orf19.2541 | Tat-D | Protein with endonuclease activity | PRO | (15, 44) |  |
| orf19.5943.1 | Stm1 | Protein of unknown function | ANTI | (34, 36) |  |
| Rny11 | Rny1 | Protein with endonuclease activity | PRO | (48)26 |  |
| Svf1 | Svf1 | Putative survival factor | ANTI | (30, 33) |  |

**Table 3.** Results from SRM and DIA comparison from the protein set that conformed the SRM method

| Protein | Hydrogen peroxide |  | Acetic acid |  |
| --- | --- | --- | --- | --- |
|  | 5 mM | 10 mM | 40 mM | 60 mM |
| Bcy1 | ✓ | ✓ | ✓ | ✓ |
| Cdc48 | ✓ | X | ✓ | ✓ |
| Cpr3 | DIA | DIA | ✓ | ✓ |
| Cyc1 | X | X | ✓ | ~ |
| Efg1 | ~ | ~ | X | ✓ |
| Mca1 | ✓ | X | ✓ | X |
| Nde1 | - | - | - | SRM |
| Nma111 | DIA | DIA | X | X |
| Nuc1 | X | X | X | X |
| orf19.4423 | - | - | SRM | SRM |
| orf19.5943.1 | X | X | X | ~ |
| orf19.643 | - | - | SRM | SRM |
| orf19.916 | - | - | SRM | SRM |
| Oye2 | ✓ | ✓ | X | X |
| Oye32 | ✓ | ~ | X | X |
| Pet9 | DIA | DIA | DIA | DIA |
| Por1 | ✓ | X | X | ✓ |
| Ras1 | ✓ | X | ~ | X |
| Rsm23 | DIA | DIA | ✓ | DIA |
| Svf1 | ✓ | ✓ | ✓ | ~ |
| Tdh3 | ~ | ~ | ✓ | X |
| Tma19 | X | X | ✓ | ✓ |

“✓” Same result in DIA and SRM
“X” Different result in DIA and SRM
“~” not significant change in one of the strategies but same ratio tendency
“-” unable to quantify by DIA or SRM
“SRM” only quantified by SRM
“DIA” only quantified by DIA
The comparison was performed taking into account significant p-values <0.05 but without a threshold in ratio (ratio>0 or ratio<0).

**Table 4.**
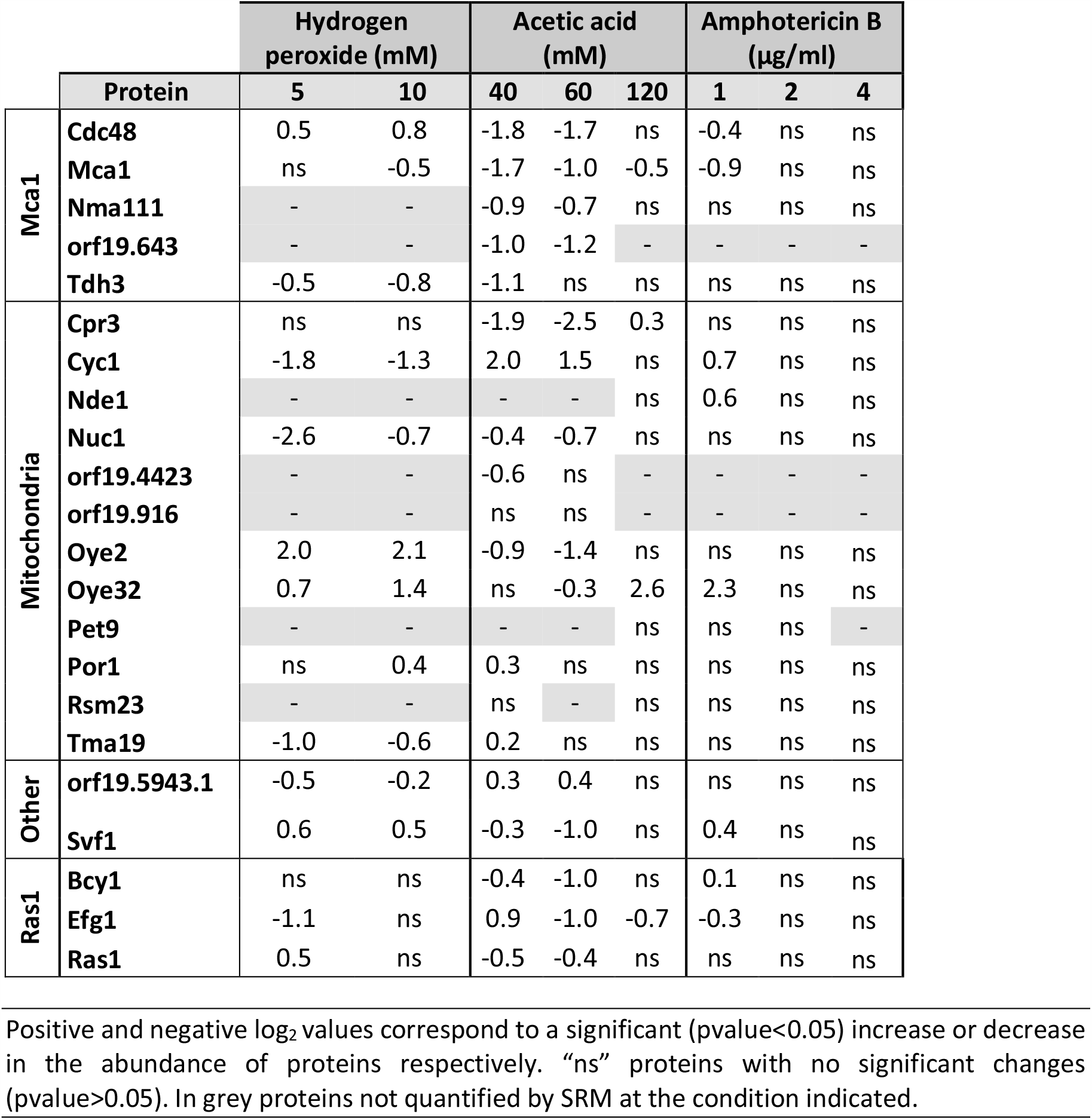
Results from SRM quantitation of proteins involved in regulated cell death

|  |  | Hydrogen peroxide (mM) |  | Acetic acid (mM) |  |  | Amphotericin B (µg/ml) |  |  |
| --- | --- | --- | --- | --- | --- | --- | --- | --- | --- |
| Protein |  | 5 | 10 | 40 | 60 | 120 | 1 | 2 | 4 |
| Mca1 | Cdc48 | 0.5 | 0.8 | -1.8 | -1.7 | ns | -0.4 | ns | ns |
|  | Mca1 | ns | -0.5 | -1.7 | -1.0 | -0.5 | -0.9 | ns | ns |
|  | Nma111 | - | - | -0.9 | -0.7 | ns | ns | ns | ns |
|  | orf19.643 | - | - | -1.0 | -1.2 | - | - | - | - |
|  | Tdh3 | -0.5 | -0.8 | -1.1 | ns | ns | ns | ns | ns |
| Mitochondria | Cpr3 | ns | ns | -1.9 | -2.5 | 0.3 | ns | ns | ns |
|  | Cyc1 | -1.8 | -1.3 | 2.0 | 1.5 | ns | 0.7 | ns | ns |
|  | Nde1 | - | - | - | - | ns | 0.6 | ns | ns |
|  | Nuc1 | -2.6 | -0.7 | -0.4 | -0.7 | ns | ns | ns | ns |
|  | orf19.4423 | - | - | -0.6 | ns | - | - | - | - |
|  | orf19.916 | - | - | ns | ns | - | - | - | - |
|  | Oye2 | 2.0 | 2.1 | -0.9 | -1.4 | ns | ns | ns | ns |
|  | Oye32 | 0.7 | 1.4 | ns | -0.3 | 2.6 | 2.3 | ns | ns |
|  | Pet9 | - | - | - | - | ns | ns | ns | - |
|  | Por1 | ns | 0.4 | 0.3 | ns | ns | ns | ns | ns |
|  | Rsm23 | - | - | ns | - | ns | ns | ns | ns |
|  | Tma19 | -1.0 | -0.6 | 0.2 | ns | ns | ns | ns | ns |
| Other | orf19.5943.1 | -0.5 | -0.2 | 0.3 | 0.4 | ns | ns | ns | ns |
|  | Svf1 | 0.6 | 0.5 | -0.3 | -1.0 | ns | 0.4 | ns | ns |
| Ras1 | Bcy1 | ns | ns | -0.4 | -1.0 | ns | 0.1 | ns | ns |
|  | Efg1 | -1.1 | ns | 0.9 | -1.0 | -0.7 | -0.3 | ns | ns |
|  | Ras1 | 0.5 | ns | -0.5 | -0.4 | ns | ns | ns | ns |

Positive and negative log<sub>2</sub> values correspond to a significant (pvalue<0.05) increase or decrease in the abundance of proteins respectively. "ns" proteins with no significant changes (pvalue>0.05). In grey proteins not quantified by SRM at the condition indicated.

We use the SRM method created to measure the participation of these proteins in *C. albicans* exposed to 1, 2, and 4 µg/ml of AMB and 120 mM acetic acid, conditions previously described as apoptotic (16, 22). We previously confirmed that all these conditions led to a loss of viability but not to a high percentage of loss of membrane permeability (always less than 1%, except less than 8% for 4 µg/ml AMB) (Fig. 7). The SRM analysis revealed changes in the abundance of proteins at 1 µg/ml AMB and 120 mM AA. There were no significant changes at higher doses of AMB (2 and 4 µg/ml) (Table 4).

**Figure 7.**
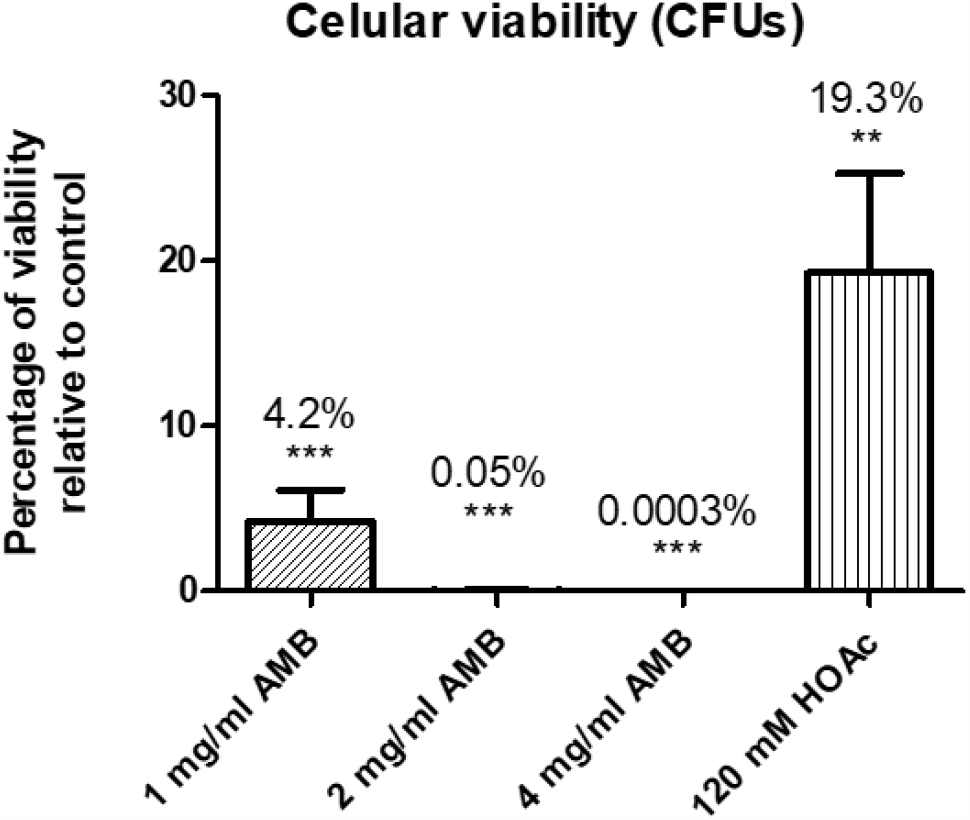
Percentage of viable cells after treatments with several concentrations of Amphotericin B or 120 mM acetic acid compared to control samples. A significant change is indicated as *pvalue<0.05, **pvalue<0.01, ***pvalue<0.001 (paired t-test). Results represent the mean of at least three biological replicates.

Interestingly, the protein Oye32 increased in abundance upon treatment with 120 mM acetic acid and 1 µg/ml AMB (Fig. 8) as it did in response to treatment with 5 mM and 10 mM hydrogen peroxide. In view of these results, we wished to determine whether these increases in Oye32 abundance were also correlated with an apoptotic state. Apoptosis occurred in 46% and 67% of cells treated with 120 mM acetic acid and with 1 µg/ml AMB, respectively (Fig. 9). These results place Oye32 in the spotlight as a common marker of *C. albicans* apoptosis induced by multiple stressors.

**Figure 8.**
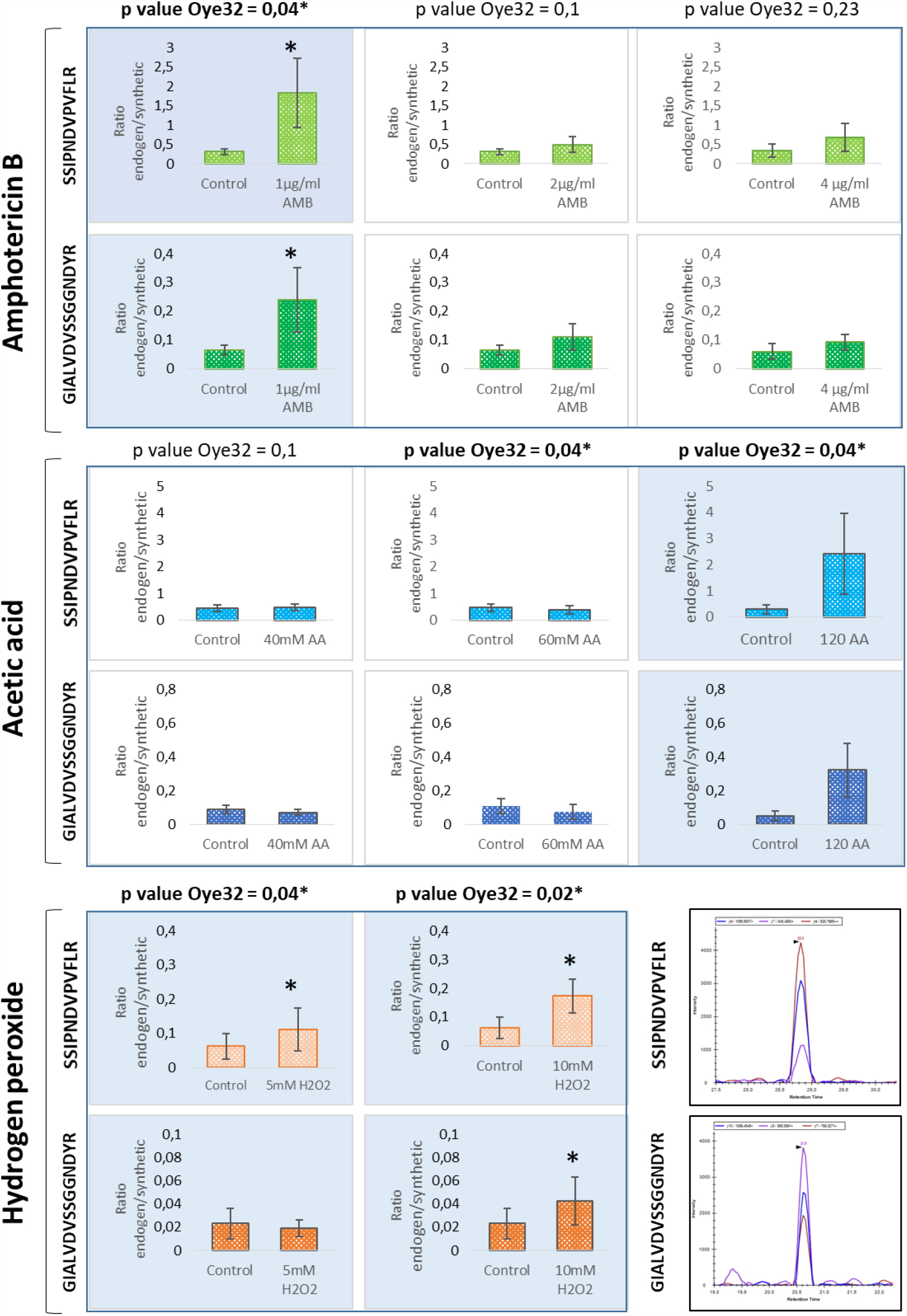
Quantification of peptides from Oye32 protein by SRM after the treatment with AMB, acetic acid or hydrogen peroxide. Blue shade in graphs indicate the conditions were apoptosis was demonstrated by PS exposure and an increase in Oye32 protein was observed (*). In the lower left corner is showed a representative chromatogram from the two peptides of Oye32 quantified by SRM.

**Figure 9.**
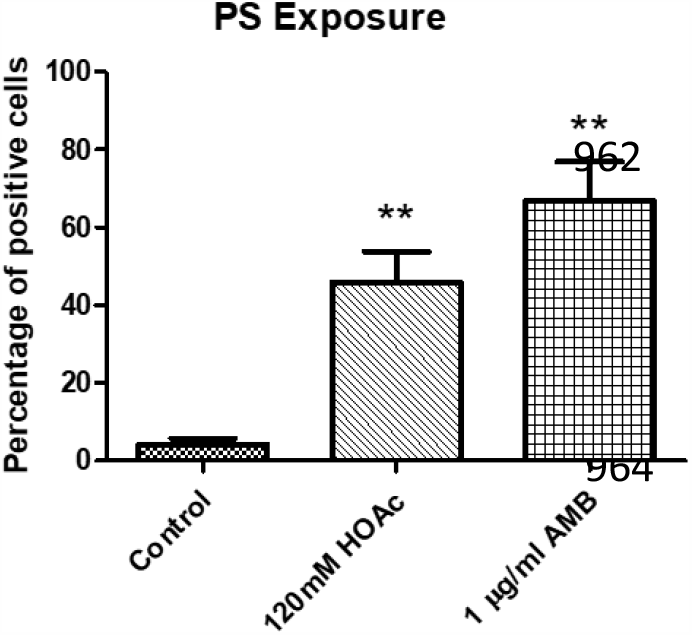
Percentage of cells showing PS exposure after treatments with 1 µg/ml Amphotericin B or 120 mM acetic acid compared to control samples. A significant change is indicated as *pvalue<0.05, **pvalue<0.01, ***pvalue<0.001 (paired t-test). Results represent the mean of at least three biological replicates.

## DISCUSSION

We have enlarged the map of the *C. albicans* global proteomic response to hydrogen peroxide and acetic acid as representative stressors in the host. The high-throughput DIA-MS proteomic analysis is one of the most extensive characterizations of *C. albicans* proteome remodeling in response to these conditions to date (52, 55). The analysis also provides a global vision of the proteome upon apoptosis induced by hydrogen peroxide in *C. albicans*. Both concentrations of hydrogen peroxide assayed (5 mM and 10 mM) stimulated apoptotic cell death in *C. albicans*, as demonstrated by PS exposure and accompanied by an increase in ROS production and caspase-like enzymatic activity. In this context, the proteomic results revealed increases in the abundance of proteins involved in antioxidant defense systems, proteasome-mediated catabolism, and protein folding, indicating an active response of the fungus against the agent. The proteasome plays an essential role in the removal of oxidatively damaged proteins, so the results are consistent with the environmental insult promoted by hydrogen peroxide. Moreover, proteins involved in protein folding were also increased in abundance, consistent with a need to repair oxidatively damaged proteins. Intriguingly, a higher dose of hydrogen peroxide was more effective in inducing apoptosis and resulted in increased levels of fewer proteins belonging to the highlighted biological processes. This result suggested a failure in the proteomic response of cells upon more severe treatment. The minor antioxidant response observed after exposure to 10 mM hydrogen peroxide, including key elements of this response, could be what induced apoptosis in a larger percentage of cells (Fig. 10). Our proteomic results reflect the same scheme of oxidative damaged proposed by Costa *et al*. in *S. cerevisiae* (56). According to Costa and colleagues, an increase in ROS will lead to the oxidation of proteins, which can be repaired by antioxidant systems or degraded by the proteasome if irreversibly damaged. Extensively oxidized proteins can form aggregates that cannot be degraded, impairing 20S proteasome and mitochondrial function. Our results demonstrate a reduction in proteasome activity in 10 mM compared with 5 mM hydrogen peroxide. Furthermore some mitochondrial proteins from the respiratory chain such as Cox4, Cox5, Cox6, Cox8, Cox9 and Qcr7 were also less abundant after the treatment (56).

**Figure 10.**
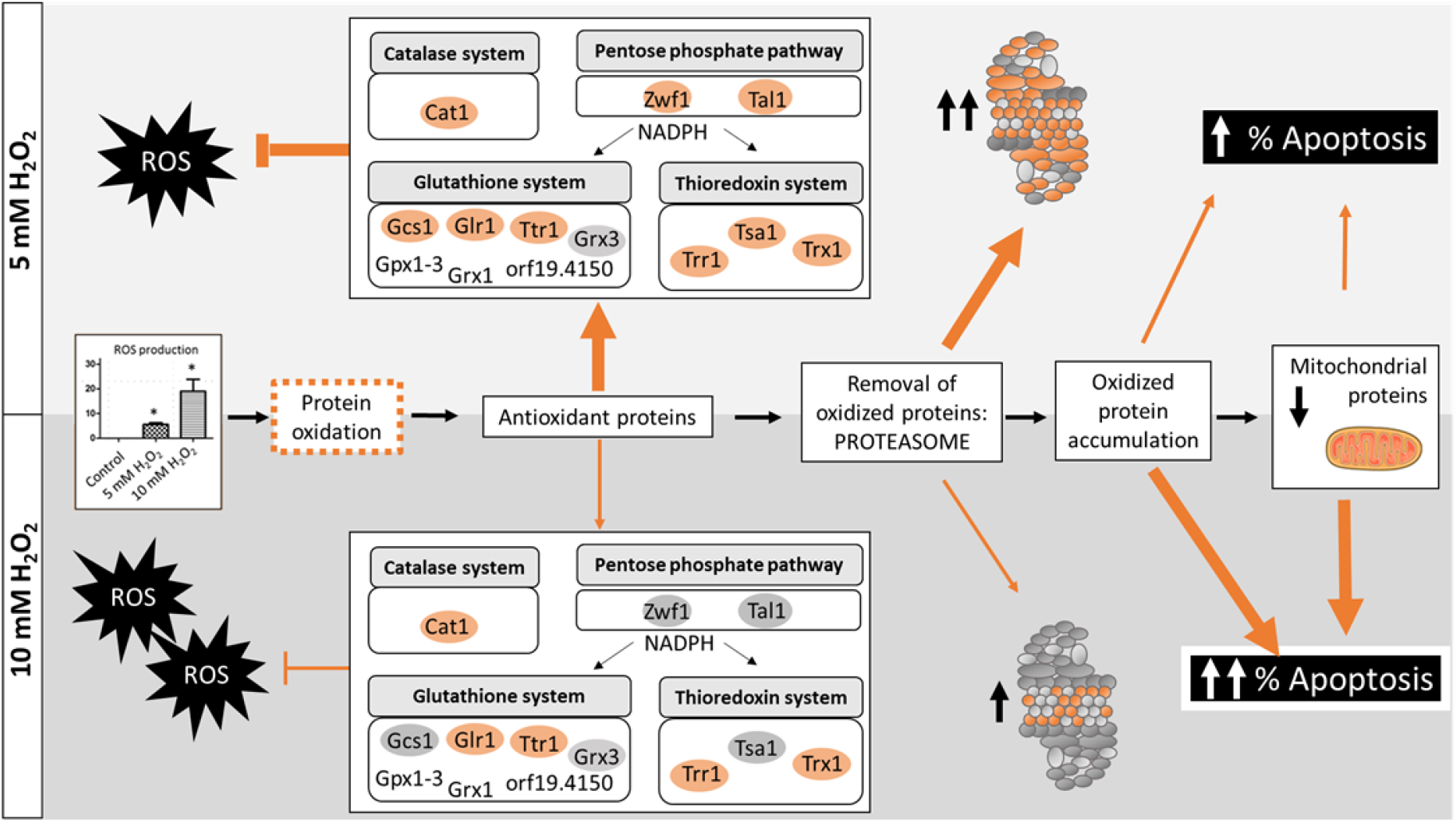
Global overview of *C. albicans* response to hydrogen peroxide according to the results from this work. Oxidative damage in proteins started with the increase in ROS production which triggered an antioxidant response with the increase of proteins from main detoxification systems (catalase, glutathione and thioredoxin). The increase of antioxidant proteins was less remarkable than at the higher concentration of hydrogen peroxide (10 mM) leading to the accumulation of oxidized proteins. This effect join to the less activity of the proteasome observed at 10 mM than 5 mM hydrogen peroxide conducted to mitochondrial damage and an increase in apoptosis percentage. The thickness of lines indicate the activation of each process. Adapted from Costa *et al* 2007.

In addition, our proteomic study uncovered the relevance of Prn1, a protein with unknown function in *C. albicans* that, according to our results, protects the cell from oxidative stress. This is supported by the increase in sensitivity to oxidative stress of a Prn1 deletion mutant. Prn1 is similar to the pirins, proteins that have been related to cellular stress and whose overexpression has been linked to apoptosis in multiple kingdoms (57, 58). For instance, in *Streptomyces ambofaciens* a protein similar to the pirins has been associated with resistance to oxidative stress (59). Our results are the first evidence of a role for Prn1 in the response to oxidative stress in *C. albicans*. This protein lacks a homologue in *S. cerevisiae*, suggesting that it may be specific to pathogenic fungi. This crucial finding highlights the usefulness of global proteomic analysis in revealing the functions of unknown proteins.

Global proteomic analysis of *C. albicans* cells subjected to acetic acid stress provided detailed information about the main cellular components that are altered in response to a fungistatic agent. Few large-scale studies have focused on the response of *C. albicans* to acetic acid; some of them have analyzed the transcriptome exclusively, and most of them have been performed on *S. cerevisiae* for its interest to industry (60-63). Our results showed a dramatic decrease in the abundance of numerous proteins involved in amino acid biosynthesis, oxidative stress, protein folding, the proteasome, actin polymerization, and translation, correlated with arrested cell growth. Our results agree with previous results from *S. cerevisiae* demonstrating a block in amino acid biosynthesis, which could be favorable for economy in energy (63, 64). Despite the increase in ROS production of cells upon acetic acid insult, the abundance of many proteins related to oxidative stress protection declined, a phenomenon previously described in *S. cerevisiae* (62, 65). Some authors have suggested that an increase in ROS upon acetic acid exposure could somehow aid the cell’s response (64).

Surprisingly, ribosomal subunit proteins were notably more abundant upon acetic acid treatment, in contrast with the general decreased observed in proteomic pattern. This might be explained by the decreased abundance of the proteasome subunits and consequent loss of proteasome activity, which could be lead to an accumulation of ribosomal proteins usually produced in excess by cells (66). Ribosomal proteins are among the most prominent ubiquitin-conjugated species that accumulate upon proteasome inhibition. Consequently, the decline in proteasome activity caused by acetic acid could explain the increase in the abundance of up to 30 proteins from the large and small ribosomal subunits. In addition, despite the presence of large amounts of ribosomal subunit proteins, the observed reduction in levels of proteins that are crucial for accurate rRNA processing could prevent the efficient formation of mature ribosomes and thus interfere with translation (67).

Some key cellular components responded to various agents in opposite manners. While the increase in proteasome subunits after exposure to hydrogen peroxide would be protective, the decrease in the abundance of proteasome proteins upon acetic acid treatment and consequent inhibition of proteasome activity could, presumably, cause ribosomal subunits to accumulate.

This study included a detailed proteomic characterization of the behavior of subunits from the core and regulatory particles of the proteasome. The functional validation of the proteomic data linked the detected changes in the amounts of proteins to changes in proteasome activity. The proteasome has been ascribed a dual role in regulated cell death (68). Some proteasome inhibitors are attracting interest for their potential therapeutic use (69). For instance, bortezomib, a proteasome inhibitor used in cancer therapies, enhances the activity of fluconazole in antifungal treatment (70). In light of our results, the slight increase in proteasome activation would affect the cell to counterbalance the oxidative stress leading to apoptotic cell death.

Unlike global proteomics, a targeted proteomic approach can focus on relevant processes within the cell such as apoptosis. Similar results obtained by the two methods support the validity of the observed changes. The different results from the two methods observed for some proteins could be explained by the use of different peptides in the quantification. In these cases, the change in the abundance should be considered with caution and based on quantotypic peptides in the SRM method, where the number of peptides used is limited (71). The SRM method could be improved by using peptides that were efficiently detected and quantified in the DIA approach. Moreover, some proteins that could only be quantified using the SRM method demonstrated that the integration of the two approaches contributes to novel results.

Cells treated with hydrogen peroxide exhibited an increase in relevant proteins such as Oye2 and Oye32, oxidoreductases described in *S. cerevisiae* for their role in apoptosis (41)(Table 4). Furthermore, the observed increase in Ras1 abundance is in accord with a prior study of apoptosis induced by hydrogen peroxide (16). The increase in Svf1 (a putative survival factor), which is crucial for the survival of cells under oxidative stress, revealed the effort of cells to overcome the stress. The increase in Cdc48 was closely related to proteasome activity and the decrease in mitochondrial proteins such as Cyc1, Nuc1, and Tma19 also reflected the alteration of these cellular components previously unmasked by the global proteomic analysis using DIA-MS. Overall, these results support the relevance of these proteins to apoptotic cell death induced by 5 mM and 10 mM hydrogen peroxide, and reinforce the use of the SRM method to evaluate the roles of proteins in apoptosis.

Despite the absence of an apoptotic phenotype (PS exposure) in cells treated with acetic acid, we used the SRM method to study the behavior of the selected proteins under stress. The SRM results were consistent with the findings of our DIA-MS global proteomics analysis: the decline in the levels of most proteins in the SRM panel was evidence for the arrested state of cells treated with acetic acid.

The targeted proteomic approach revealed a consistent increase in the abundance of Oye32 in cells subjected to various stresses that can induce apoptosis (5 mM and 10 mM hydrogen peroxide, 120 mM acetic acid, and 1 µg/ml AMB) suggesting this protein as a possible apoptotic biomarker.

The SRM method designed in this study is a straightforward proteomic approach to monitoring apoptotic proteins that might be targets for anti-fungal therapies in several circumstances. The existence of regulated cell death in fungal pathogens could be usefully exploited to develop novel antifungal therapies (6, 72).

## MATERIALS AND METHODS

### Fungal strains and culture conditions

*Candida albicans* wild-type strain SC5314 was used for the phenotypic and proteomic analysis performed. For hydrogen peroxide susceptibility assays control strain SN250 and *prn1* mutant from Noble collection were used (73). Yeast cells were maintained at 30°C on YPD medium containing 1% yeast extract, 2% peptone and 2% glucose with rotatory shaken (180 rpm). Before exposure to agents cells were grown until exponential phase was reached (optical density 1 ± 0.2). Then acetic acid, hydrogen peroxide or Amphotericin B (AMB) were added to the desired concentration and incubated for 200 min at 30°C with rotatory shaken. For the phenotypic and proteomic assays after treatment with hydrogen peroxide a final concentration of 5 mM and 10 mM were selected (Sigma-Aldrich). For susceptibility assays *C. albicans* strains growth at optical density of 1 were exposed to 80 and 100 mM hydrogen peroxide for 0, 2, 5, 10 min. Four microliters of yeast suspension were spotted on YPD agar plates and incubated for 48 h at 30°C.

The acetic acid concentrations used in the experiments were 40, 60 and 120 mM (PanReac, AppliChem). For AMB assays a stock solution (1 mg/ml) was prepared in DMSO (Amphotericin B, 85%, Acros OrganicsTM, Fisher). AMB was added to an exponential culture at final concentrations of 1, 2 and 4 µg/ml for 200 min at 30°C in rotatory shaken.

### Viability assays

To determine the percentage of viable cells after suspend the treatment, cells were collected and washed three times with PBS. Optical densities (620 nm) were measured to normalize the amount of cells and correlate with the number of colony forming units (CFUs) growth on plates after 48 h at 30°C.

### Loss of cell membrane integrity

Propidium iodide was used to evaluate the loss of selective permeability of the membrane after treatments. Cells were stained with 5 µl of propidium iodide (50 µg/ml) for 5 min at room temperature. Percentage of positive cells was calculated by observation on a fluorescence microscope.

### Externalization of phosphatidylserine

Phosphatidylserine (PS) externalization was evaluated by staining protoplasted cells with Annexin V-FITC (Takara). Protoplast were generated as previously described (74). Briefly, *C. albicans* cells were incubated with 0.5 ml of 50 mM K_2_HPO_4_, 5 mM EDTA, and 50 mM DTT (adjusted to pH 7.2) at 30°C for 30 min to promote spheroplast formation. After that, 0.5 mL of solution containing 50 mM KH_2_PO_4_, 40 mM 2-mercaptoethanol, 0.15 mg/ml zymolyase 20T and 20 µl of glusulase in 2.4 M sorbitol (pH 7.2) was added and incubated for 30 min at 30°C and 80 rpm. Protoplast were stained following manufactured instructions from ApoAlertTM annexin V-FITC kit (Takara) in modified annexin binding buffer containing 1.2M sorbitol. Flow cytometry was used to determine the percentage of cells annexin positive indicating early and late apoptosis.

### ROS production

*C. albicans* cells from control and treatments were washed thrice in cold PBS and stained with 5 µg/ml dihydrorhodamine 123 (DHR-123, Sigma) for ROS production 30 min before finish the experiment (74). The percentage of ROS positive cells was evaluated with a fluorescence microscope by counting at least 100 cells from at least 3 biological replicates.

### Caspase like enzymatic activity

The increase in caspase like enzymatic activity was evaluated by using a staining solution containing (FITC)-VAD-FMK (CaspACE™, Promega) at a final concentration of 10 µM/ml (74). Cells were incubated for 20 min at 37°C, washed twice with PBS and counted in a fluorescence microscope.

### Preparation of soluble protein extracts and peptide digestion for mass spectrometry

Cells from control and treated samples were harvested and washed thrice in cold PBS. Then, cells were resuspended in lysis buffer (50 mM Tris-HCl pH 7.5, 1 mM EDTA, 150 mM NaCl, 1 mM DTT, 0.5 mM PMSF, and 10 % of a mix of protease inhibitors (Pierce TM) and disrupted by adding glass beads (0.4-0.6 mm diameter) in a Fast-Prep system (Bio101, Savant) applying 5 cycles of 21 sec. Cell extracts were separated from glass beads by centrifugation and the supernatant was collected and cleared by centrifugation at 13000 rpm for 15 min at 4°C. Protein concentration was measured by using Bradford assay, and 50 µg of protein extracts were prepared for digestion.

Peptide digestion for SRM and DIA was performed using 50 µg of cytoplasmic extracts that were denatured with 8 M Urea in 100 mM ammonium bicarbonate and reduced with Tris (2-carboxyethyl) phosphine 5 mM for 30 min at 37°C. After that samples were alkylated with 10 mM iodoacetamide for 45 min in dark. Finally samples were digested by trypsin (1/100, w/w, Promega) for 16 h at 37°C. Peptides were purified using reverse phase C18 columns. Peptide quantitation was performed using BCA (Pierce Quantitative Colorimetric Peptide Assay) or Qubit (Thermo Scientific) system.

### Data Dependent Acquisition (DDA) and Data Independent Acquisition (DIA) set up

All global proteomic analysis were performed on a Q Exactive Plus mass spectrometer (Thermo Scientific) connected to an EASY-nLC 1000 ultra-high-performance liquid chromatography system (Thermo Scientific). Peptides were separated on an EASY-Spray column with a linear gradient of 5-30% acetonitrile (ACN) in aqueous 0.1% formic acid, during 60- or 120-minutes for DDA or DIA, respectively. Settings for DDA and DIA analysis were the same as previously described by Malmström (75). Shortly, acquired spectra were analysed using the search engine X! Tandem (Jackhammer, 2013.06.15) against Assembly 21 (A21-s02-m09-r10) from Candida Genome Database (CGD, (76)) or synthetic retention time peptide sequences with reverse decoys (77). X!Tandem parameters were fixed to 1 missed peptide cleavages and the following modifications: fixed carbamidomethilation of cysteine and the N-terminal variable modifications: acetylation, cyclisation of S-carbamoylmethylcysteine, and pyroglutamic acid formation of glutamic acid and glutamine. The precursor and fragment mass tolerance was set to 20 ppm and 50 ppm, respectively. Peptide spectrum matches were filtered to FDR>1%.

The spectral library was generated on DDA mode from 52 samples from *C. albicans* upon different conditions (unpublished data deposited at PRIDE repository, project accession: PXD020195). The assay library was assembled using Fraggle that interprets and averages MS2 spectra. After, we used Franklin to perform a multilevel FDR calculation and finally, the assay library was generated with Tramler (78).

*C. albicans* spectral library generated was used to extract DIA data from each condition using DIANA algorithm v2.0.0 with FDR correction set to 1% for identification and quantification (79). For protein quantification, the top3 integrated peptide ion intensities from MS2 spectra were considered (80).

### DIA MS data analysis and GO Term enrichment

For relative quantitation, protein intensities were previously normalized within each sample. Proteins identified in at least 3 biological replicates were considered for quantitative analysis. Statistical analysis by paired t-test from at least 3 biological replicates and were performed. Significant changes in the abundance of proteins between control and treatments were considered when pval < 0.05 and log_2_ fold change more than or less than −0.5 (paired t-test from at least 3 biological replicates).

Significant proteins were selected to perform GO term analysis using GO Term Finder tool from CGD and Genecodis (76, 81). Representative biological processes and cellular component altered after treatments were selected according to p value and number of proteins included.

Heatmaps and volcano plots were performed using RStudio (v1.0.143) using ggplot2 and pheatmap.

### SRM design and quantitation

#### 1. Peptide/transition selection and validation

Targeted proteomic assays were performed according to the method previously reported (82). A set of 32 proteins involved in yeast apoptosis were selected for the targeted proteomic analysis based on previous works published. A total of 63 proteotypic peptides (peptide uniquely associated with the protein of interest) from the selected proteins were designated for SRM relative quantification based on information from PeptideAtlas (83). The same peptides were purchased as synthetic heavy labeled peptides (JPT Peptide Technologies) and used in unpurified form. To confirm peptide identities and select the best three transitions for SRM method, fragment ion spectra from each peptide included in yeast matrix were acquired by MS2 analysis in a QTRAP 5500 instrument (AB/SCIEX). The data was also used to know retention times and optimize the method in schedule mode. An adjusted amount of each synthetic peptide was added to a final solution containing all peptides according to their signal intensities. The ratio of synthetic peptides added to the peptide sample was 1:5.

#### 2. SRM mass spectrometer configuration

Digested samples with synthetic peptides included were injected into a nHPLC (Eksigent nano LC 1D plus) where peptides were concentrated in a trap column (Eksigente nanoLC trap) for 5 minutes at a flow of 2 µl/min of 2% ACN, 0.1% AF before their separation in a 15 cm analytical column (Eksigent nanoLC column 2C18-CL). Peptide elution was achieved by a 5-35% ACN gradient for 30 min at 300 nl/min. Both trap and analytical columns were heated at 50 °C to get more constant retention times between samples. At least three biological replicates of each condition were analyzed, with at least 2 technical replicates of each one. All analysis were performed in an AB Sciex QTRAP 5500 mass spectrometer.

#### 3. SRM data analysis

The raw files containing SRM data were analyzed with Skyline software (v 4.2.0.19009), the peak selection in the chromatograms of each peptide was manually curated (84). Transitions showing some interference in peak area were excluded. The intensity area of each peak was automatically calculated by the software considering the value as the ratio endogenous/synthetic precursor. The statistical tool implemented in Skyline software reported the significant difference of endogenous peptide between samples (t-test<0.05).

### Determination of enzymatic activity of the proteasome

For the measurement of proteasome activity a fluorometric assay kit (Proteasome 20S Activity Assay kit) that measures the chymotrypsin-like protease activity was used following manufactured instructions (Sigma-Aldrich). A total of 100 µg of protein extracts were incubated with LLVY-R110 substrate provided by the kit for 2 h at 30°C in dark. The green fluorescent signal generated by the cleavage of LLVY-R110 by proteasome was measured in a BMG FLUOstar Galaxy equipment (λex= 480–500 nm/λem= 520–530 nm). Data from three technical replicates and three biological replicates measured as fluorescence arbitrary units were used for paired t-test analysis (p value <0.05).

## ACKNOWLEDGEMENTS

This study was supported by BIO2015-65147-R and RTI2018-094004-B-100 from Spanish Ministry of Science and Innovation, InGEMICS-CM B2017/BMD3691 from the Comunidad de Madrid, Spanish Network for the Research in Infectious Diseases (REIPI RD16/0016/0011) and PRB3 (PT17/0019/0012) from the ISCIII. InGEMICS-CM, REIPI and PRB3 are financed jointly by European Development Regional Fund ERDF “A way to achieve Europe”. The proteomic analyses were performed in the laboratory of Johan Malmström and in the Proteomics facility of Complutense University of Madrid a member of ProteoRed-ISCIII network. These results are lined up with the Human Infectious Diseases HPP initiative from the Human Proteome Project (HID-HPP). A. Amador-García was a recipient of a fellowship from Ministry of Science and Innovation (FPI) and from the Comunidad de Madrid (Youth Employment Initiative, European Commission).

## DATA AVAILABILITY

The data set from this paper has been deposited in the ProteomeXchange Consortium via the PRIDE partner repository with the data set identifier PXD020283.

## SUPPLEMENTAL LEGENDS

**S1:** GO enrichment analysis from proteins more or less abundant after treatment with 5 mM or 10 mM hydrogen peroxide

**S2**: Proteins from cell wall and involved in ATP synthesis that decrease their abundance after treatment with hydrogen peroxide

**S3**: GO enrichment analysis from proteins more or less abundant after treatment with 40 mM or 60 mM acetic acid

**S4**: SRM method

